# Inferring the shape of global epistasis

**DOI:** 10.1101/278630

**Authors:** Jakub Otwinowski, David M. McCandlish, Joshua B. Plotkin

## Abstract

Genotype-phenotype relationships are notoriously complicated. Idiosyncratic interactions between specific combinations of mutations occur, and are difficult to predict. Yet it is increasingly clear that many interactions can be understood in terms of *global epistasis*. That is, mutations may act additively on some underlying, unobserved trait, and this trait is then transformed via a nonlinear function to the observed phenotype as a result of subsequent biophysical and cellular processes. Here we infer the shape of such global epistasis in three proteins, based on published high-throughput mutagenesis data. To do so, we develop a maximum-likelihood inference procedure using a flexible family of monotonic nonlinear functions spanned by an I-spline basis. Our analysis uncovers dramatic nonlinearities in all three proteins; in some proteins a model with global epistasis accounts for virtually all the measured variation, whereas in others we find substantial local epistasis as well. This method allows us to test hypotheses about the form of global epistasis and to distinguish variance components attributable to global epistasis, local epistasis, and measurement error.

## Introduction

The mapping from genetic sequence to biological phenotype in large part determines the course of evolution. Non-additive interactions between sites, called epistasis, can either accelerate or severely constrain the pace of adaptation. Due to the high-dimensionality of sequence space, studies of evolution on epistatic genotype-phenotype maps, or fitness landscapes, have historically been limited to mathematical models, such as NK landscapes [1, 2], or computational models of simple biophysical phenotypes, such as RNA folding [3, 4].

Recently, high-throughput sequencing has made it possible to measure genotype-phenotype maps for proteins [5] and across genomes [6]. While many thousands of genotype-phenotype pairs can now be assayed in a single experiment, this is still only a minuscule fraction of all possible sequences, and so the sampled genotypes typically take the form of a scattered cloud centered on the wild-type. Statistical models are therefore required to derive biological insight from these sparsely sampled genotype-phenotype maps. One approach has been to fit models that include terms for each pairwise interaction between genetic sites [7, 8]. Such models can sometimes predict protein contacts or mutational effects from protein sequence alignments [9], but they do a poor job at capturing a complete picture of the genotype-phenotype map – so that their predictions far from the observed data are often wildly inaccurate [10, 11].

Models of a genotype-phenotype map that contain terms for every possible pairwise interaction seem reasonable if we believe that epistasis arises primarily through the idiosyncratic effects of particular pairs of mutants – for instance, pairs of residues that contact each other in a folded protein. While there is abundant biochemical evidence for epistasis of this type, geneticists have long suspected that a substantial fraction of observed epistasis is due not to specific pairwise interactions, but rather to inherent nonlinearities in molecular phenotypes, cellular fitness, organismal physiology, and reproductive success. Apparent pairwise interactions may be caused by mutations that contribute additively to some underlying trait, combined with a nonlinear relationship between the underlying trait and the measured phenotype, or fitness.

A classic argument for this form of epistasis is the physiological theory of dominance [12], which held that dominance arises from the inherently nonlinear relationship between gene activity or dosage and measured phenotype (e.g., saturation phenomena in metabolic networks [13]). Another classical example arose in the attempt to reconcile apparent contradictions between patterns of genetic polymorphism and the tolerable degree of genetic load, by positing models of nonlinear selection—for instance truncation selection—operating on an underlying additive trait such as the number of deleterious mutations or heterozygous genes [14, 15, 16, 17, 18]. Following these classic examples, research into the shape of nonlinear selection on an observed quantitative trait using either quadratic [19], or more general [20] forms became a major interest of evolutionary quantitative genetics [21]. And contemporary studies on the fitness effects of mutations often incorporate epistasis based on thermodynamic arguments, where the probability of transcription factor binding [22] or protein folding [23, 24, 25] is expressed as a nonlinear function of a binding or folding energy that is additive across sites.

Today, the idea that epistatic interactions are the result of a nonlinear mapping from an underlying additive trait has been called “uni-dimensional” [26, 27], “non-specific” [28] or “global” [29] epistasis, and several heuristic techniques have been proposed to infer epistasis of this form [30, 31, 32, 33].

Here we present a maximum-likelihood framework for inferring models of global epistasis, and we apply this framework to several large genotype-phenotype maps of proteins derived from deep mutational scanning (DMS) experiments. Under the assumption that the observed phenotype is a monotonic, nonlinear function of some unobserved additive trait, we find the best-fit model of global epistasis, test hypotheses about the form of this nonlinear relationship, infer the additive coefficients for the unobserved trait together with their confidence intervals, and estimate the extent of additional epistasis due to specific interactions between sites. Our approach shares much of the simplicity of additive models, but it can capture far more of the phenotypic variation using only a handful of additional parameters.

### A model of global epistasis

Statistical models can infer genotype-phenotype maps from data by assuming some form of its underlying structure. The starting point of our analysis is a non-epistatic model. Given a sequence of amino acids *a*_*i*_ for sites *i* = 1 to *L* and an associated measured phenotype *y*, the non-epistatic model is

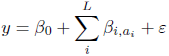

where the noise term *ε* represents deviations between model and data. The additive effects *β*_*i,α*_ can be visualized as a matrix of effects for each position and amino acid with *β*_*i,ai*_ = 0 if there is no substitution (i.e.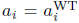), so that the wild-type phenotype is modeled by the constant term *β*_0_. With many pairs of measured sequences and phenotypes the additive effects *β*_*i,α*_ can be estimated by linear regression. Non-epistatic models often explain much of the variance in genotype-phenotype maps, but there is usually a significant portion of unexplained variance that cannot be accounted for, given the known magnitudes of measurement errors [34].

One obvious way to incorporate epistasis into this additive model is by including explicit terms for interactions between each pair of sites, and possibly higher-order interactions as well [7, 8]. By contrast, we develop a model and inference procedure for a different form of epistasis, motivated by examining the deviations between empirical data sets and their best-fit non-epistatic models. Such deviations often show a smooth nonlinearity, as seen in Fig. S1A for protein G. This observed nonlinearity suggests a global coupling between all sites that determines the observed phenotype. This idea can be formalized as a semi-parametric model

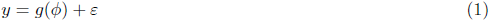

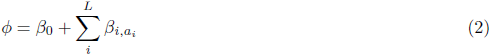

where *ϕ* is an inferred, but unobserved, additive trait that depends on the genetic sequence, and *g*(*ϕ*) is an arbitrary nonlinear function that represents the shape of global epistasis relating the additive trait to the measured data. We refer to the coefficients *β*_*i,α*_ in this model as the additive effects, even though the full model is nonlinear. The assumption underlying this model is that mutations affect the measured phenotype only via a single unobserved additive trait. To model the function *g*(*ϕ*) we choose a flexible parametric family of monotonic functions, I-splines [35], in order to avoid fitting an arbitrarily complex model and to reduce inference to standard problems in maximum likelihood. With sufficient data, the additive effects *β*_*i,α*_ and the shape of global epistasis *g*(*ϕ*) can be inferred simultaneously.

We also estimate the amount of epistasis that is not captured by the global nonlinearity in our model. When independent estimates of the measurement error for each sequence are available, as is often the case, we can infer how much of the remaining variation in the measured phenotype is due to other forms of epistasis. We refer to this last component of variation in the measured phenotype as house-of-cards (HOC) epistasis, because in our statistical framework we model the phenotypic contribution of this epistasis as an independent random draw, for each genotype, from a Gaussian distribution – akin to a fully uncorrelated house-of-cards HOC model [36]. In particular, we estimate the HOC epistasis component, 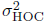, by setting the total variance of our per-sequence Gaussian likelihood to be 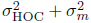, where 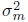is the independent estimate of phenotype measurement error (see Methods).

We infer the model described above by maximum likelihood. Doing so allows us compare models of different complexity by means of a likelihood ratio test via parametric bootstrap (detailed in *Methods*). This approach allows us to ask whether there is statistical support for certain features, but not others, in the shape of global epistasis, and to assess the uncertainty in our estimates of both global epistasis and the coefficients describing the impact of mutations on the underlying additive trait.

## Global epistasis in protein GB1

As a first application of our framework, we characterized global epistasis in the IgG-binding domain of protein G (GB1), a model system of protein folding and stability (fig. 1), using data from a study by Olson et al. [37].

**Figure 1:**
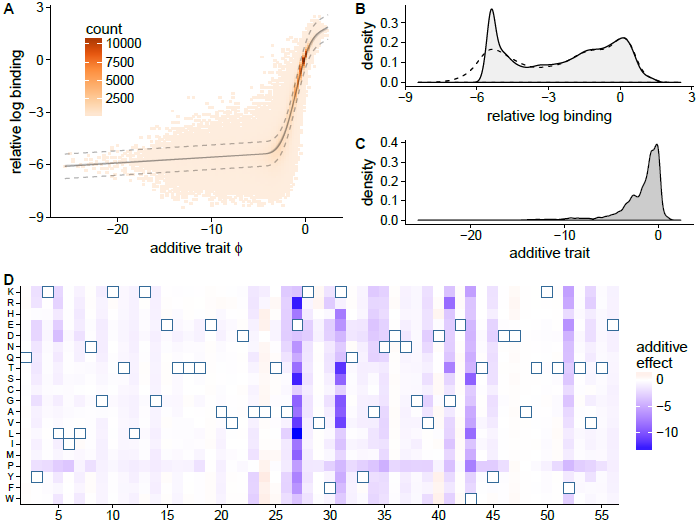
A) Global epistasis as a nonlinear function of an intermediate additive trait (eq. 1) for binding of protein GB1 to an immunoglobulin fragment (IgGFC) [37]. Solid black line indicates the nonlinear function *g*(*ϕ*) (95% confidence interval in light gray). Based on the inferred magnitude of non-global epistasis, we estimate that the true phenotypes for 95% of genotypes lie between the two dotted lines (2*σ* _HOC_ = 0.7). Colors are a histogram of observed binding. B) The distribution of binding values are bimodally distributed (measured dashed line, inferred solid line), while the distribution of inferred additive phenotypes is unimodal (C). D) The impact of each possible amino acid substitution on the underlying additive trait. Outlined squares are the wild-type amino acid. Additive effects and the underlying trait are scaled so that the wild-type has trait value 0 and the average mutation changes the trait by distance 1 (see *Methods*)

In particular, Olson et al. targeted 55 sites for mutation and measured the binding to an immunoglobulin fragment (IgGFC) for all single amino acid substitutions (1045), and 95% of all double substitutions (509,693) [37]. We find that for GB1 our model of global epistasis outperforms a purely non-epistatic model (*p* < 0.0003, likelihood ratio test, Tab. S1, see methods for statistical methodology), and it substantially reduces the extent of unmodeled epistasis (σ _HOC_ = 0.35, reduced from σ _HOC_ = 0.56 for the non-epistatic model) with a 10-fold cross-validated *r*^2^ = 93.5%, compared to *r*^2^ = 86.2% for the non-epistatic model.

Looking first at the shape of global epistasis inferred by our model (fig. 1A), we see that it includes both diminishing and increasing returns depending on the value of the underlying additive trait. In particular, our inferred non-linear function has a negative second derivative in the region around the wild-type (negative in 95% of bootstraps for *-*0.97 *< ϕ <* 1.72) indicating a pattern of diminishing returns epistasis, but the slope remains strictly positive at high trait values (*p <* 0.003, tab. S1) suggesting that further increases in binding affinity outside the range of observed data are possible. For deleterious mutations, we observe saturation with decreasing additive trait values, as indicated by a positive second derivative (i.e., increasing returns; positive in 95% of bootstraps for *-*4.27 *< ϕ < -*1.04). We find significant support for a nonzero slope on the left-hand side (*p <* 0.003, tab. S1), but our estimate of this slope is extremely small, with an expected change of only 0.0037 (CI 0.0034 to 0.0046) per mutation, in units of relative log binding. This slope is very small compared to the overall range of binding scores observed (from around −5 to +2) and it is consistent with control experiments by [37] that established a lower bound on the binding score in their assay. More generally, our estimates of the shape of global epistasis are extremely precise, with an average 95% confidence interval of width 0.13 (see *Methods*) over the full range of the observed binding affinities (gray region in fig. 1A).

For the effects on the unobserved additive trait (fig. 1D), we find 234 beneficial and 774 deleterious mutations with 95% CIs that exclude zero, out of a total of 1045 mutations (fig. S1D). These additive effects are similar to what would be inferred under the purely additive model, but they are more widely distributed for deleterious mutations (fig. S1B). Interestingly, the distribution of inferred additive trait is unimodal whereas the observed distribution of binding affinities is bimodal, with one peak around the wild-type affinity and another at low-binding (fig. 1B, C). This apparent contradiction is resolved by the form of global epistasis, which maps all sequences with low values of the additive trait to similar values for the observed phenotype, creating a second mode.

Taken together, our model for the underlying additive trait and for the nonlinear function *ϕ* provides a simple explanation for the observed binding data. The model accounts for essentially all the variation in binding measurements beyond what would be expected under measurement noise alone. In particular, root mean squared error in our predictions is 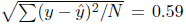 whereas even under a perfect model would expect a root mean squared error of 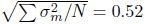. the due to the Poisson variation in the counts used to calculate the binding scores The extent of these errors is very small compared to the overall range of phenotypes observed, and it constitutes a sequence-function relationship that is dominated by global epistasis with little room for epistasis of other kinds. This can be understood visually by examining fig. 1A, where we predict that the true phenotypes for 95% of genotypes fall between the two dashed lines.

## Complex epistasis in GB1

While our model captures the form of global epistasis and quantifies the extent of remaining HOC epistasis, deviations from our model can provide evidence for one remaining type of epistasis, which we call complex epistasis. This form of epistasis arises from interactions between specific combinations of sites. Unlike global epistasis, which depends in a nonlinear way on a single additive trait, or HOC epistasis, which consists of independent Guassian effects associated with each genotype, complex epistasis can be attributed to nonlinear interactions between specific sites on the measured phenotype.

To investigate complex epistasis we compared the predictions of our model, fitted to data from Olson et al. [37], to an independent dataset from a followup experiment by [38]. Wu et al. measured the binding of GB1 variants mutated to all possible combinations of amino acids at four sites specifically chosen because they were suspected to exhibit idiosyncratic epistatic interactions.

On the whole, our results confirm that the pattern of global epistasis inferred using shallow mutagenesis across the whole domain persists at a deeper level of mutagenesis for these four sites. In particular, the predictions of the global epistasis model, fitted to Olsen et. al data, are close to the mean binding curve measured by Wu et al., across the entire range of the inferred additive trait (fig. S2A). However, the width of the deviations from our binding predictions is much broader than the variation predicted by the error model fit to the Olsen et. al. experiment (fig. 1A). Some of this difference can be attributed to the much larger measurement noise in the [38] dataset 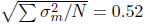 Olsen et al. versus 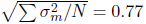 Wu et al.) due to the lower read-depth in Wu et al. However, the deviations from the model are also skewed – so that around 13 % of phenotypes measured by Wu et. al have significantly higher binding than predicted, even after the difference in measurement noise is taken into account (fig. S2B). We interpret this skew as evidence of positive complex epistasis among the specific sites mutagenized by Wu et. al.

To infer which combinations of sites and mutations in GB1 exhibit complex epistasis we marginalize the distribution of deviations from our model predictions for each pair of substitutions. Pairs of mutations with large mean squared deviations (fig. S2C) indicate systematic deviation from the global epistasis model. For example, the pair of mutations 41L/54G shows the largest deviation in our analysis, and indeed it is known to be associated with structural changes in GB1 [37]. By contrast, the pair 41Q/54P exhibits a similar magnitude of pairwise epistasis calculated directly from binding data and yet, in our analysis, the deviations from the model are small, indicating this pairwise epistasis is primarily caused by global epistasis. Accounting for global epistasis can therefore be instrumental in identifying the specific sites that have idiosyncratic, complex interactions for a measured phenotype.

## Global epistasis in GFP

The GB1 dataset of Olsen et. al contains only single and double mutants, which covers a limited cloud of sequence space near the wild-type protein sequence. To characterize a genotype-phenotype maps more broadly we applied our model of global epistasis to a DMS study that measured the fluorescence of 51,715 variants of a green fluorescent protein (avGFP), including up to 11 mutations per sequence and 3.7 mutations on average [31].

Applied to the GFP data, our model of global epistasis infers a sharp threshold that delineates fluorescing from non-fluorescing proteins, with remarkably few outliers (true positive rate 0.9966, true negative rate 0.9787, where we define positive sequences to have log relative fluorescence of −1.25 or greater, fig. 2). Below the threshold, where the fluorescence is at background levels, the trait-fluorescence relation is almost flat (slope 0.00032 log fluorescence per mutation, 95% CI 0.00022 to 0.00041); while above the threshold the global epistasis function is increasing (likelihood ratio test *p <* 0.003, tab. S1, slope 0.020 log fluorescence per mutation, 95% CI 0.019 to 0.021). In other words, mutations observed to have no effect below the threshold may have a substantial effect in a different genetic background.

**Figure 2:**
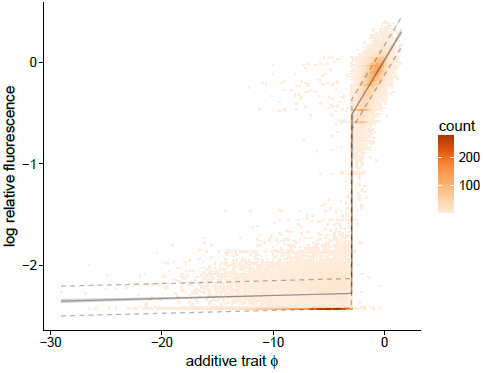
The shape of global epistasis in a DMS study of GFP [31] shows a sharp threshold in the additive trait. Below the threshold there is low fluorescence and above the threshold fluorescence is high and the slope of the non-linear mapping is positive (likelihood ratio test *p <* 0.003). Data consists of 51,715 protein variants, with 3.7 mutations on average. HOC epistasis has magnitude *σ* _HOC_ = 0.073, and 10-fold cross validated *r*^2^ = 0.931 0.003. *g*(*ϕ*) indicated by black line with gray shadow indicating the 95% bootstrap confidence interval. Our analysis suggests that the true values for 95% of genotypes will lie between the dashed lines. The underlying additive trait is scaled so that the wild-type has trait value 0 and the mean absolute magnitude mutation changes the trait by distance 1 (see *Methods*)

Comparing the epistatic and non-epistatic models (fig. S3A) shows that including global epistasis im-*CV* proves the fit substantially (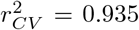compared for 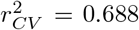for the non-epistatic model), and the*CV* inferred flourescent variance attributable to HOC epistasis is much smaller after accounting for global epistasis (*σ* _HOC_ = 0.073 versus *σ* _HOC_ = 0.31). Whereas for GB1 we found very small confidence intervals both for *g*(*ϕ*) and for the additive effects of mutations on the underlying trait (e.g. fig. S1D), for GFP we find that the bootstrapped 95% confidence intervals for *g*(*ϕ*) are very small (mean 0.18), but that the infered additive coefficients (fig. S3C) show much greater uncertainty than observed for GB1. In particular, we find that only 62% of the additive coefficients have confidence intervals excluding zero (141 positive and 1131 negative coefficients out of 1810) total, and the average width of these confidence intervals was equal to 0.91 times the magnitude of an average mutation. These results show the importance of quantifying uncertainty, and they suggest that although the GFP experiment had sufficient enough power to infer the form of global epistasis, it was under-powered with respect to inferring the effects of individual mutations on the unobserved, additive trait.

Overall, our results indicate that the sequence-function relationship for GFP can be adequately understood in terms of a sharp threshold imposed on an underlying additive trait. Below this threshold, fluorescence is zero, and and above this threshold fluorescence is essentially additive in the effects of mutations. More specifically, our model suggests that 95% of genotypes will fall between the two dashed lines in fig. 2, and the error in our predictions is little more than would be expected under measurement noise alone (mean squared error in our predictions is 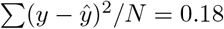, whereas we would expect a root mean squared error of 0.16n due to measurement error alone, see Methods). Thus, like GB1, we find that global epistasis plays a dominant role in the GFP genotype-phenotype map, with little need to invoke other forms of epistasis to explain the observed data. These results are essentially consistent with the results of the neural network approach used by [31] in their original analysis (based on a neural network architecture with a sharp bottleneck to represent the latent trait), but our approach provides a simpler, easier to understand picture of the sequence-function relationship together with better quantification of uncertainty.

## Genotype-by-environment interactions in *β*-lactamase

Deep mutational scanning studies often measure phenotypes across a library of variants in several different conditions – such as antibiotic activity at several concentrations of antibiotic. While testing in multiple conditions can in principle provide more insight into the function of a protein and should provide more robust results than measurements in a single condition, it also introduces a problem of interpretability. In particular, it becomes difficult to summarize the effects of any given mutation across the experimental conditions. Here we address this problem by extending our model to analyze genotype-by-environment interactions. We do this by assuming that the genotype and environmental condition each contribute additively to the latent underlying trait *ϕ*, which then determines the observed phenotype via a nonlinear function. While not universally applicable, such models make biological sense in cases where both the genotype and environment can act on some latent underlying trait. For instance, it is reasonable to assume that the action of an antibiotic depends on its “local” concentration in the cellular or extra-cellular environment, which can be modulated either through changes to the activity of an enzyme that degrades the antibiotic or by changes to the overall concentration of antibiotic in the medium.

Here we apply this approach to a study of *β*-lactamase, an enzyme that degrades *β*-lactam antibiotics, which has been a model system for deep mutational scanning [39, 40, 41] and protein evolution [42, 43, 44, 45]. Stiffler et al. [46] measured the effects of 4,997 single mutants of TEM-1 *β*-lactamase in five different concentrations of the antibiotic ampicillin. They quantified *β*-lactamase activity by measuring growth rate of each mutant in each condition. Our model takes these five growth rate measurements and synthesizes them into a single score capturing the activity of that particular genotype, together with a nonlinear function capturing the mapping from the latent trait (e.g. effective local concentration of antibiotic) to the observed growth rate.

The results of fitting this global-epistatic model are shown in fig. 3, where the x-axis displays the activity of each mutant (i.e. the impact of that mutant on the underlying trait), and the environmental conditions determine a series of curves, one for each condition. Due to our assumption that the environment and genotype interact additively to determine the underlying trait, these curves are simply translations of one another, and each curve produces a prediction of the growth rate of each mutant for one environment. Overall, we find the model produces a very good fit to the data (cross-validated *r*^2^ = 0.92), far better than a purely additive model (*p <* 0.0003, tab. S1). The performance is also comparable to the biophysically based mechanistic model originally fit by [46] which uses approximately five thousand more parameters.

**Figure 3:**
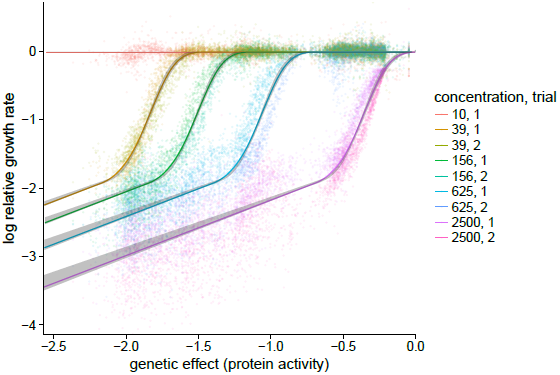
Gene-by-environment interactions in a DMS study of *β*-lactamase [46]. Data consists of 4,997 single mutants measuring growth rate under different concentrations of antibiotic (ampicillin) and two replicates [46]. Log relative growth rate is plotted against the inferred protein activity for each substitution; the mapping between protein activity for log relative growth rate for each antibiotic concentration is given by the colored curves and each such curve is surrounded by a gray region giving its 95% confidence interval. Growth rate measurements were made using two biological replicates, shown as two slightly different hues for each antibiotic concentration. Cross-validated *r*^2^ = 0.923 ± 0.002.

The global-epistasis model provides a score for each mutation’s additive effect on the underlying trait. We again normalize so that the wild-type has a score equal to zero and the mean absolute effect of a mutation on the additive trait equals 1. We infer that the single mutation effects (fig. S5) are largely deleterious: 3295 out of 3312 single mutations have CI below zero, and none has CI above zero (fig. S6B), consistent with the observation that no mutant displays a growth rate consistently higher than wild-type.

The inferred activity scores are mapped to growth rates via a nonlinear function that shifts depending on the antibiotic concentration. This function shows first increasing and then decreasing returns (for instance in the highest antibiotic concentration, the second derivative has positive 95% CI for *-*0.61 *< ϕ < -*0.38 and negative CI for −0.33 < *ϕ* < −0.09, where *ϕ* is the genetic effect on the underlying trait), and it ultimately saturates with zero slope at high concentrations (*p* = 0.39, tab. S1). These results indicate a threshold-like phenomenon where, for any given genotype, increasing the antibiotic concentration up to a certain point produces no growth defect, whereas further increases in antibiotic produce rapidly declining growth rates. Interestingly, while the slope of the nonlinear function becomes shallower for low activity genotypes (slope 0.090 fitness change/mutation on left hand side, CI 0.073-0.095), it remains significantly greater than 0 (*p <* .0003, tab. S1). This result may have clinical relevance: although growth rates experience a sudden decline with increasing antibiotic concentration, increasing the concentration beyond the point of first inhibition is still capable of producing a further decrease in bacterial growth rate. Finally we tested the hypothesis that the environmental coefficients were a linear function of the log antibiotic concentration versus a more general model that fit an arbitrary coefficient for each condition. We found strong evidence against the linear model (*p <* 0.003, tab. S1), consistent with the visual pattern of increasingly severe antibiotic effects at higher concentrations (fig. 3).

The simplicity of our model and together with its graphical representation provides insights into the design and behavior of this experiment. For instance, in (fig. 3) we have colored the two biological replicates in each condition slightly different colors. The plot shows that the replicates are not completely overlapping, which indicates biological variation between the replicates; and indeed we can reject our basic model in favor of a model with a different environmental coefficient for each antibiotic concentration in each replicate. Another fine scale pattern can be observed in the confidence intervals for the additive effects (Fig. S6B), which widen for genotypes that have WT-like growth rates at one antibiotic concentration and then very low growth rates in the next concentration. This suggests a finer grid of antibiotic concentrations would be optimal for future experiments.

## Additive traits and biophysical quantities

Since we infer an underlying additive trait when fitting a global epistasis model, it is natural to ask whether the underlying trait corresponds to some known biophysical quantity. While this question will surely depend upon the details of the assay used in a given deep mutagenesis scanning experiment – e.g., whether the assay is measuring binding of a protein to a substrate, enzymatic activity, etc., thermodynamic stability for a protein’s native conformation is almost always and important phenotype, and there is a consistent thread in the literature suggesting that the phenotypic effects of most mutations are mediated via their effects on thermodynamic stability and the non-linear relationship between free energy of folding and probability of being folded [23, 24, 47, 25, 48, 49]. Moreover, low-throughput measurements of mutational effects on thermodynamic stability (∆∆*G*) for mutations relative to wild-type are typically additive [50, 51], and relatively conserved over evolutionary time [52]. Thus, under the hypothesis that most mutations affect phenotype via their energetic effects of thermal stability, the inferred additive coefficients under our model should be highly correlated with the stability effects of these same mutations.

To test this hypothesis, we calculated the correlation between our additive effects and previously published estimates of the stability effects of these mutations. For protein G, we do not find a correlation between these measured stability effects and our inferred effects on the underlying additive phenotype (fig S1E, *p* = 0.14), a result consistent with the original authors’ observation that the mutational effects on the binding phenotype of single mutants around the wild-type sequence were not correlated with their effects on protein stability [37, 53]. Similarly for *β*-lactamase, independent measurements of ∆∆*G* are not significantly correlated with our inferred additive effects (figs. S6A, *p* = 0.08). For GFP, on the other hand, the inferred additive trait is negatively correlated (*ρ* = *-*0.62, *p* = 2 × 10^−16^) with computational predictions by [31] of energetic effects for single mutations on stability, as would be predicted if the fitness effects for most mutations were mediated through their effects on folding stability. Thus while the good fit of our our model is consistent with the hypothesis that measured phenotypic effects are mostly determined by a non-linear function of the free energy of folding [23, 24, 47, 25, 48, 49], the inconsistent relationship between our inferred additive effects and the estimated stability effects of mutations provides at best equivocal support for the widely-held hypothesis that free energy of folding is the underlying trait.

## Discussion

We have proposed a maximum-likelihood framework for inferring genetic interactions from high-throughput mutagenesis experiments. Our key assumption is that genetic interactions arise from a global nonlinear mapping between an unobserved, additive trait and the measured phenotype. Modeling this nonlinear mapping by a flexible class of monotonically increasing functions, we simultaneously infer the form of the nonlinear relationship and the contributions of each possible mutation to the underlying additive trait. Analyzing three well-studied proteins, we have shown that this form of global epistasis can provide a remarkably good fit while providing an easy-to-understand summary the underlying sequence-function relationship. Our approach also provides a principled statistical framework for testing hypotheses and quantifying uncertainty.

Understanding genotype-phenotype relationships provides a substantial challenge due to the enormous size and large dimensionality of the space of possible genotypes. An ideal model of this relationship would exhibit comprehensible behavior, make accurate predictions, and give mechanistic insight into the underlying biology. In practice, however, models must make trade-offs between these competing demands. Our strategy is based on the gambit that, despite abundant evidence for complex genetic interactions between specific sites, a simple-to-understand model of global epistasis can account for most phenotypic variation. The resulting models can be summarized visually by a pair of graphics: a heat map of coefficients showing the effects of each possible single amino acid substitution on an underlying additive trait, and a curve showing the shape of the nonlinear relationship between the additive trait and observed phenotype. Because we assume that this nonlinear relationship is monotonically increasing with increasing values of the underlying trait, the resulting model is single-peaked, and it shows no sign epistasis [54]. In particular, the sign of the effect of any particular mutation is constant across backgrounds, and the magnitude of the effect depends only on the current trait rather than on the details of the genetic sequence [29]. This simplicity contrasts with the highly complex, rugged landscapes typically inferred by fitting a model with all possible pairwise interactions between sites,where assessing qualitative features of the landscape from the fitted coefficients can be extremely challenging. For instance, finding the global maximum in a pairwise interaction fit is an NP-complete problem [55], whereas the optimal genotype can be read off directly from the heat map of additive coefficients in our model of global epistasis.

Our results show that sometimes a simple model of global epistasis can provide accurate predictions that account for virtually all the observed experimental variation. Because of the high dimensionality of protein sequence space, even a completely additive model will typically have several thousand parameters. By augmenting an additive model with just a handful of additional parameters, which control the nonlinear function, we can produce a dramatically better fit and capture the majority of observed variation with many hundreds times fewer parameters than in pairwise models (e.g. *∼* 500k pairwise interactions for GB1). The large dimensionality of genotypic space also puts a premium on quantifying uncertainty. Perhaps surprisingly,we find that the inferred confidence intervals for the form of global epistasis are much tighter than the confidence intervals for the additive effects of individual mutations, e.g. in our re-analysis of GFP [31], suggesting that robust measurements of global epistasis will be possible even from limited mutagenesis data. The global epistasis model also makes reasonable predictions for genotypes far from those sampled in the mutagenesis assay, unlike the qualitatively incorrect out-of-sample predictions exhibited by pair-wise models[10, 11]. Extrapolation can be easily understood under the global epistasis framework, because it is based on the biologically plausible assumption that the physiological factors responsible for saturation or potentiation operate consistently across genetic backgrounds. Furthermore, our global epistasis model is additive outside the observed range of phenotypes and so it provides highly controlled, conservative behavior when extrapolating to extreme phenotypes.

While the global epistasis model is readily comprehensible, and it is sometimes sufficient to capture the major features of the genotype-phenotype relationship, it is unclear whether the model provides an unambiguous mechanistic interpretation in terms of molecular biology and biophysics. If most epistasis in proteins is indeed due to essentially additive effects on the free energy of folding, that are then converted by a nonlinear transformation between folding energy and the probability of folding [23, 24, 47, 25, 48], then our model should fit well and the theory of stability mediated epistasis would provide a mechanistic basis for the observed additive trait. On the other hand, the observation that our model fits well is not alone sufficient to infer the existence of a mechanistically meaningful underlying additive trait. For example, if the phenotype is a saturating function of several underlying traits (e.g. [56, 37, 53]) the additive trait inferred under our model may correspond to an amalgamation of multiple mechanistic traits.

The interpretation of the global epistasis model as the simplest possible epistatic extension of an additive model, together with its capacity to capture a large proportion of the observed epistasis, suggests that it should be used as a standard first analysis of mutagenesis assay data, instead of an additive fit. More expressive models, capable of capturing complex epistasis, can then be used to describe the idiosyncratic interactions between the small set of sites whose interactions deviate substantially from the large-scale patterns identified by the global-epistasis fit. Moreover, differences in experimental protocol can be viewed as changes in the nonlinear mapping from the additive trait to observed phenotype, and so we expect the inferred additive coefficients to be more consistent and reproduceable across laboratories and experiments than the raw measurements. Indeed for TEM-1, we find that our inferred coefficients are more highly correlated with previous measurements of TEM-1 activity [40] then the measured growth rates in any single antibiotic concentration (*r*^2^ = 0.96 for coefficients, 0.06, 0.74, 0.90, 0.91, 0.79 in individual conditions).

A subtle, but important, technical note concerns our assumption that the nonlinear global epistasis func-tion *g*(*ϕ*) is a monotonic function spanned by an I-spline basis. *Some* restriction on the form of *g*(*ϕ*) is essential because a model allowing *g*(*ϕ*) could exactly match the measured phenotypes and would therefore provide no useful simplification. In our case, we restrict the function *g*(*ϕ*) to be monotonic for comprehensibility—even a quadratic mapping can produce extremely complex patterns of epistasis, see [57]). Although other families of monotonically increasing functions are certainly possible (e.g. [58] or [59], who use very similar methods to show there is no global nonlinearity in antibody binding energies) we have found that a 4-element I-spline basis is sufficient for the datasets explored here, and adds only a handful of additional parameters relative to an additive model. In contrast, recent heuristic approaches to model global epistasis essentially suffer from imposing too strong or too weak constraints on the nonlinear function *g*(*ϕ*) (for instance, [31, 32] use a neural network approach that lacks easy comprehensibility, whereas [33] assume that the nonlinear function is a power transformation, which is comprehensible but too constrained to capture physiologically realistic patterns of saturation).

The intuition that complicated behavior in a high-dimensional space can often be summarized by learning a nonlinear function of a latent additive trait has a long history in biology—as detailed in the introduction. And techniques for inferring such relationships have been rediscovered multiple times across a breadth of scientific disciplines (e.g. single-index models in econometrics [60], projection pursuit regression in statistics [61], linear-nonlinear models in neuroscience [62]). Here we have shown that these ideas can be productively applied to modeling genotype-phenotype relationships in high-throughput mutagenesis data. While the form of epistasis is very simple and easy-to-understand, our model nonetheless captures the major features of genetic and genotype-by-environment interaction for GB1, GFP and TEM-1 beta-lactamase. We suggest that it provides a natural first model to apply to any high-throughput mutagenesis dataset.

## Methods

### Preprocessing of genotype-phenotype data

As input our procedure requires pairs of genotype-phenotype measurements and optional estimates of measurement error and environmental condition.

For the GB1 datasets [37, 38], we defined a functional score and variance from the sequence counts before and after selection based on a poisson approximation

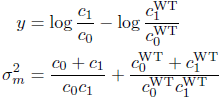

where *c*_0_ and *c*_1_ are the counts before and after selection, respectively, and both include a pseudocount of 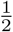 [63]. The score is defined relative to the wild-type sequence, using counts 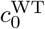 and 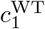.

For the GFP dataset [31], we used the fluorescence measurement (minus the wild-type fluorescence) and variance as provided.

For the *β*-lactamase dataset [46], trials 1 and 2 were combined, with given functional measurements relative to wild-type (and no estimates of measurement error). The wild-type sequences were not given, and therefore we added them to our data with value equal to zero for each trial and antibiotic concentration. Mutations were excluded that showed small effects relative to the distribution of neutral variants (absolute difference from wild-type less than 0.25 under all concentration of antibiotic) because the additive genetic effect of mutations that show no fitness defect across all conditions is not well-defined under our best-fit model (see below).

### Inference of non-epistatic model

The additive trait *ϕ* is defined as in eq. 2 and the goal is to infer the parameters. The sequence is encoded as a set of zeros and ones representing each possible substitution from the wild-type (dummy coding). We define a Gaussian likelihood for each observation *y* (representing binding, fluorescence, etc.) with mean equal to *ϕ*, and variance that is the sum of the measurement error 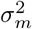 for each sequence and the HOC epistasis 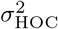.This results in 19*L* + 1 parameters for the trait and the HOC epistasis variance. We find the parameters that maximize the sum of log-likelihoods per observation with a standard gradient-based algorithm, L-BFGS (NLopt library), with parameters initialized from the coefficients of a non-epistatic model fit to the data using ordinary least squares.

### Inference of global epistasis

Our model of global epistasis is that the measured phenotype is a nonlinear function, *g*, of an additive trait, *ϕ*. For the nonlinearity, we implemented a flexible family of 3rd order monotonic I-splines with *k* evenly spaced knots [35]. We found that four knots were sufficient for each analysis and had sufficient flexibility without overfitting. The nonlinear function is a linear combination of the I-spline basis functions plus a constant, *g* = *c*_*α*_+*∑*_*m*_ *α*_*m*_ *Im*, with *k* + 1 non-negative *α*_*m*_ coefficients. These functions are only defined within the knot boundaries, so we extend them to linearly extrapolate beyond the knot boundaries. We defined a likelihood for each observation *y* analogous to the non-epistatic model, with a mean equal to *g*(*ϕ*),and variance 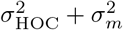, and maximized this likelihood using the optimization algorithm described above.

The optimization is sensitive to the choice of initial parameters, and we used the following procedure to HOC choose those parameters. We fit a non-epistatic model to get *β* _*i,α*_ and 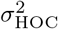 and then rescale them such that the trait has a range 0 ≤ *ϕ* ≤ 1We choose evenly spaced knots between zero and one, and then find *α*_*m*_ that produce a linear function *g* with a separate optimization holding other parameters fixed.

One subtly in the inference procedure is that the latent additive trait is only defined up to an affine transformation. This occurs because given an affine transformation of the underlying phenotype *A*(*ϕ*), a compensatory change in the global epistasis can give identical predictions: *g*(*ϕ*) = *g*^′^ (*ϕ*^′^) = *g*(*A*^-1^ (*A*(*ϕ*))). During the inference itself we fix the affine transformation by choosing the knot positions, keeping these fixed throughout the maximization. We then post-process the inferred parameters to have a natural biological interpretation by choosing the affine transformation that sets the wild-type phenotype to 0 and makes the average absolute value of the additive coefficients equal to 1.

Finally, some datasets have only partial information on measurement errors. For example, in GFP [31] the estimated errors are based on the fluorescence variance of sequences with the same barcode. Many of the sequences appeared only once, which has no associated variance. In this case, we extended our likelihood such that if there was no associated measurement error, we set the variance of the error term for these sequences HOCto 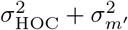 where we again estimate the parameter 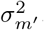.

### Separating genetic and environmental effects

We separate genetic and environmental effects with a model where the measured phenotype is a nonlinear function of an intermediate trait plus the environmental effect, *g*(*ϕ* + *ϕ*_*e*_). For a categorical environmental effect where *z* is the environmental state, *ϕ*_*e*_ = ∑*γ ζ* _*γ*_ *δ* _*γ,z*_, where *ζ* _*γ*_ is the effect of being in environment *γ* (relative to the reference environment) and *δ* _*γ,z*_ = 1 when *γ* is equal to *z* and zero otherwise. For a continuous variable *ϕ*_*e*_ = *ζz*.

As noted in the main text, our best fit nonlinear function *g* for the *β*-lactamase data had slope equal to zero,*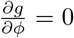*, to the right of the highest knot position. Strictly speaking, the associated coefficients *β* _*i,α*_ for any genotype that is in the flat portion of the curve for all environmental conditions become unidentifiable, as a change in the coefficient does not change the prediction. In these instances, the optimization is allowed to finish, then the *β*_*i,α*_ that are affected are decreased to be equal to the rightmost knot position.

When the data consists of single mutants in multiple environments, the cross-validation of the model requires some modification to avoid assigning all data on a given mutation to the test set, since we have no way of making predictions for mutations that have never been observed. To avoid this problem, we generate the training and testing subsets based on the environmental conditions. For each mutation that is observed under several conditions/replicates, we assign a random permutation of integers between one and the number of replicates and conditions (i.e., the 9 categories in fig. 3. These integers define the subsets used for training and testing. For each fold of cross validation, all observations assigned to a particular integer are used as the test set and the remaining observations are used as training.

### Bootstrapped confidence intervals

We estimated confidence intervals by a parametric bootstrap based on the maximum likelihood predictions. The bootstrapped data was generated as 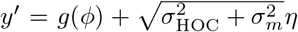, where *η* is an instance of a standard normal random variable. Because the values for the underlying trait are only defined up to an affine transformation, for each bootstrap *b*, we find parameters that linearly transform the original trait estimates into the bootstrap trait scale, *ϕ*_*b*_ = *m*_*b*_*ϕ* _ML_ + *c*_*b*_ via least squares regression, where the *ϕ*_*b*_ and *ϕ* _ML_ are vectors describing the intermediate traits in the bootstrapped and maximum likelihood models, and *m*_*b*_ and *c*_*b*_ are the inferred parameters. For calculating the confidence interval of *g*_ML_, we make a distribution based on each bootstrapped model *g*_*b*_ (*m*_*b*_*χ* + *cb*) for each value of *χ*. We choose 101 linearly spaced values for *χ* in the non-linear region (between zero and one) and 101 linearly spaced values for the each of two linearly extrapolated regions (from the knot boundaries to the maximum/minimum *ϕ*). For the confidence interval of maximum likelihood *β* _*i,α*_ we make a distribution based on *β* _*i,α,b*_ */m*_*b*_.

### Model comparison

We compared nested maximum likelihood models by a bootstrapped likelihood ratio test. First the difference in log-likelihood was computed between the simpler (null) model and the more complex (hypothesis) model. The likelihood optimization of the more complex model had initial parameters determined by the simpler (null) model. P-values were calculated by comparing to a bootstrapped log-likelihood difference distribution, with *y*^′^ generated by a parametric bootstrap from the null model as described above.

We validated this procedure by bootstrapping the p-values for the first test in table SS1. That is, for each bootstrap in the test, we calculate a p-value by means of a second (or recursive) likelihood ratio test based on the bootstrapped data and models (with 100 bootstraps). The resulting p-value distribution (fig. S7) should be uniform in the ideal case, however the resulting distribution has excess probability in the center indicating that the test is somewhat conservative.

## Supplemental Figures

**Figure S1:**
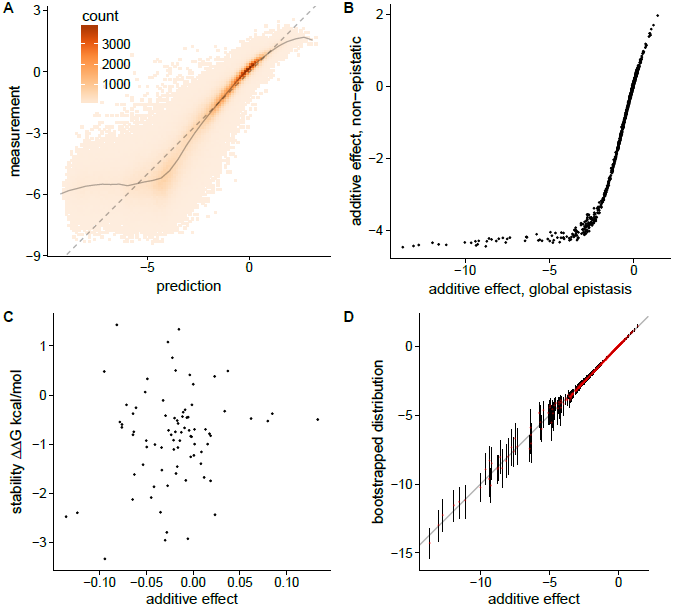
GB1 A) non-epistatic model of GB1 data shows distinct non-linearity in the predicted vs measured phenotype, *σ* = 0.314, *r*^2^ = 0.862, CV *r*^2^ = 0.862 ± 0.002. B) Inferred additive effects of non-epistatic and global epistatic models. The global epistasis model infers greater differentiation among the highly deleterious coefficients due to the non-linearity of *g*(*ϕ*). C) *β* vs independently measured ∆∆*G* for 81 substitutions (collected in [37]), correlation 0.17, p=0.13. D) Bootstrapped additive coefficients quantify the uncertainty in the parameters. Red points indicate median and lines indicate 95% confidence intervals. Out of 1045 additive effects, 234 had CI above zero, and 774 had CI below zero.

**Figure S2:**
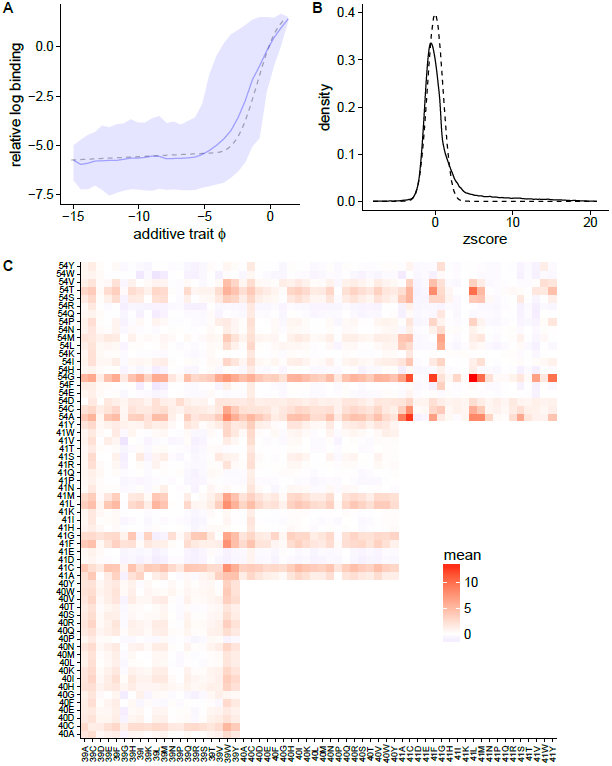
A) The global epistasis in a followup study of GB1 [38], which targeted four sites, is similar to that in the original study 1. Additive trait and binding is predicted for Wu et al. data from a model trained on Olson et al. [37]. The dashed line indicates the predicted binding, and the blue is the measured binding distribution, conditional on *ϕ*, with the solid line indicating the mean and 95% of the data is in the shaded region. B) Around 14% of the sequences in Wu et al. have much higher binding than predicted, indicating some complex epistatic effects are restoring binding. The solid line is the distribution of z-scores, that is the *m* difference between predicted and measured binding, normalized by 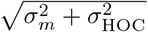 for each sequence. Dashedline is a standard normal distribution. C) Z-scores averaged over all sequences with a specific pair of mutations show how much each pair deviates from the model. The pairs 41L/54G and 41Q/54P had strong epistasis (7.1 and 6.5 respectively) [37]. While 41L/54G has a large average deviation from the global epistasis model, the pair 41Q/54P has a small deviation, indicating the pairwise epistasis may be due to global epistasis alone.

**Figure S3:**
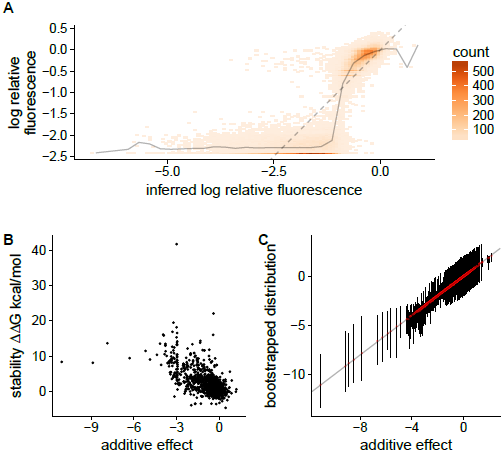
A) The non-epistatic model for GFP [31]. Colors indicate the 2D histogram of fluorescence values and the solid line is the conditional mean measured fluorescence for 30 bins. *r*^2^ = 0.71 cross-validated *r*^2^ = 0.69 ± 0.01 *σ* _epi_ = 0.31. B) Inferred additive effects and computationally determined changes in fold stability from [31]. Correlation *ρ* = −0.62, *p* = 2 × 10*-*16. C) Bootstrapped distribution of each additive effect. Lines are 95% confidence intervals and red points are medians. Out of 1810 effects, 141 have CIs above zero and 1131 have CIs below zero.

**Figure S4:**
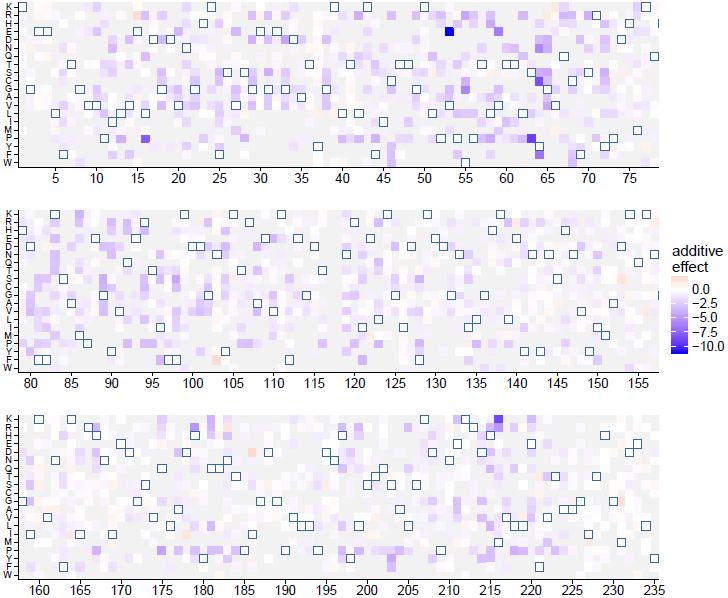
Additive effects for GFP [31]. Outlined squares are the wild-type amino acid. Gray squares denote unobserved mutations.

**Figure S5:**
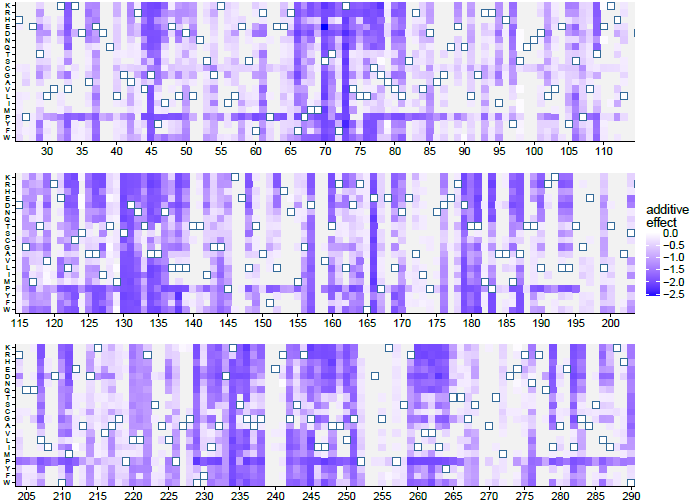
Genetic activity scores (*ϕ*) relative to the wild-type for *β*-lactamase [46]. Outlined squares are the wild-type amino acid. Gray squares denote unobserved mutations.

**Figure S6:**
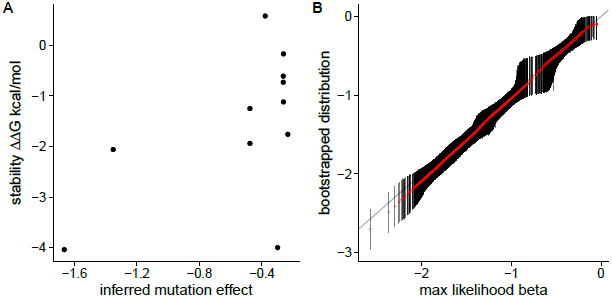
A) Measured stabilities of mutations and inferred additive trait values. Values from [64, 65, 66].B) Bootstrapped distribution of each genetic activity score. Lines are 95% confidence intervals and red points are medians. Out of 3312 effects, none have CIs above zero and 3312 have CIs below zero.

**Figure S7:**
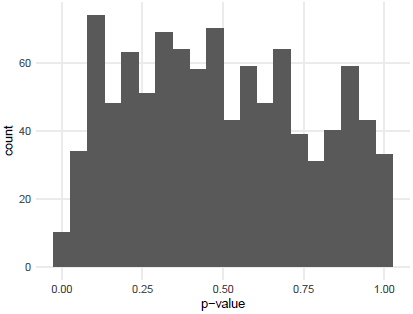
Bootstrapped p-values from bootstrapped likelihood ratio test where null is global epistasis with zero slopes at both knot boundaries for beta lactamase data, and alternative has non-negative slope on left hand side. See *Methods*.

**Table S1:** Likelihood ratio tests for various null models and their alternatives. NE is non-epistatic model, GE is global epistasis model with (true/false, true/false) indicating whether left and right side slope of *g*(*ϕ*) is constrained to zero. P-values calculated by parametric boostrap (see *Methods*). P-value is *<* 0.0003 when the change in likelihood is greater than any of the 1000 bootstrapped changes in likelihood.

| system | null | alternative | p-value |
| --- | --- | --- | --- |
| GB1 | NE | GE (false, false) | < 0.003 |
| GB1 | GE (true, false) | GE (false, false) | < 0.003 |
| GB1 | GE (false, true) | GE (false, false) | < 0.003 |
| GFP | GE (false, true) | GE (false, false) | < 0.003 |
| GFP | GE (true, false) | GE (false, false) | 0.005 |
| β-lac. | NE | GE (false, true) | < 0.003 |
| β-lac. | GE (false, false) | GE (false, true) | < 0.003 |
| β-lac. | GE (false, true) | GE (false, false) | 0.39 |
| β-lac. | continuous antibiotic effect | categorical antibiotic effect | < 0.003 |

